# Bioanalyzer chips can be used interchangeably for many analyses of DNA or RNA

**DOI:** 10.1101/039040

**Authors:** Jessica Davies, Tom Denyer, James Hadfield

## Abstract

The Agilent 2100 Bioanalyzer (Agilent Technologies, CA, USA) enables small-scale gel electrophoretic separation of nucleic acids on a microfluidic chip. Shortage of chips and excess reagents is a common issue. This report explored the compatibility of two commonly used Bioanalyzer reagents with three Bioanalyzer chip types. Microfluidic electrophoretic separation of DNA and RNA using DNA High Sensitivity and RNA 6000 Nano reagents, respectively, was successfully performed on multiple chip types, following the assay-specific protocol. For RNA quality and next-generation sequencing library size estimation, the Bioanalyzer chips tested can be used interchangeably. These findings will be valuable for any laboratory using the Agilent Bioanalyzer in a shared facility.

## Methods summary

Four RNA samples (from murine brain) and four DNA samples (RNA-seq libraries made using the Illumina TruSeq Stranded mRNA kit, Illumina Inc., San Diego, USA) were run in triplicate using RNA 6000 Nano and DNA High Sensitivity (HS) reagents, respectively, on RNA 6000 Nano, DNA HS and DNA 1000 chips. Quality and concentration of RNA samples, and concentration and size distribution of DNA samples were tested. We demonstrate that any of the Bioanalyzer chips tested can be used interchangeably with defined Bioanalyzer reagents for qualitative analysis, and can also be reasonably quantitative, provided the protocol and software method for the corresponding assay reagent kit are followed correctly.

## Main text

The Agilent 2100 Bioanalyzer performs microfluidic electrophoretic separation on microfabricated chips (1). In comparison to slab gel electrophoresis, the Bioanalyzer provides many advantages: separation is quick; minimal sample volume is required (1 μl); user exposure to hazardous materials is minimal; and the assessment of sample quantity and quality is not dependent on the user’s interpretation.

Results of nucleic acid sample separation are displayed on an electropherogram and gel-like image, generated by the Bioanalyzer 2100 Expert Software (1). These provide a visualisation of sample quality and quantity; for RNA, integrity is additionally assessed by a software algorithm which produces an RNA integrity number (RIN) (2, 3). DNA samples, such as PCR products, restriction digests, and plasmid digests, can be assessed with kits covering a vast range of product lengths. Moreover, the additional high sensitivity reagents are particularly useful for library assessment prior to next-generation sequencing.

A common problem in the laboratory is that of chip shortage with excess reagents, particularly in laboratory service environments which experience both high usage and fluctuations in the demand for different kit types. Therefore, an investigation into whether reagents can be used interchangeably with different Agilent Bioanalyzer chips would be valuable for many researchers. Anecdotal reports of using the wrong chip type have been noted previously (4). Others have demonstrated the ability to re-use chips multiple times without detrimentally affecting results (5, 6). We explored the compatibility of RNA 6000 Nano and DNA HS Bioanalyzer reagents with three chip types, following the assay-specific protocol and using the assay-specific software.

The RIN and concentration of four RNA samples measured in triplicate (R1, R2, R3, and R4) were assessed using the RNA 6000 Nano reagents and protocol, on RNA 6000 Nano, DNA HS and DNA 1000 chips. Importantly, the sticker displaying the chip layout was disregarded and the loading pattern indicated in the assay-specific protocol was used. All chips were run on the Agilent 2100 Bioanalyzer using the Eukaryotic Total RNA Nano assay. Concentration and RIN of each RNA sample were highly comparable between chips (inter-chip concentration: p = 0.96; RIN: ANOVA p = 0.13) **(see Table 1 and Fig. 1).** Intra- and inter-chip variability for RNA RIN and concentration were very similar. RIN and RNA concentration were both well within the normal variability expected of samples submitted for RNA-seq experiments.

**Table 1 legend:** RNA data. Four RNA samples were run according to the RNA 6000 Nano protocol, on each chip. RINs (upper) and sample concentration data (lower) (ng/μl) were determined. Sample R4 was excluded from the DNA HS chip, as the final three wells were used to validate ladder consistency within a chip. M = mean.

| RNA Sample | DNA 1000 |  |  |  | DNA HS |  |  |  | RNA Nano |  |  |  |
| --- | --- | --- | --- | --- | --- | --- | --- | --- | --- | --- | --- | --- |
|  | 1 | 2 | 3 | M | 1 | 2 | 3 | M | 1 | 2 | 3 | M |
| R1 | 9.9 | 9.9 | 10.0 | 9.9 | 9.2 | 10.0 | 9.7 | 9.6 | 9.9 | 8.9 | 9.7 | 9.5 |
| R2 | 9.9 | 9.9 | 9.9 | 9.9 | 10.0 | 9.0 | 10.0 | 9.7 | 9.8 | 10.0 | 9.2 | 9.5 |
| R3 | 10.0 | 10.0 | 10.0 | 10.0 | 9.5 | 9.8 | 10.0 | 9.8 | 10.0 | 10.0 | 10.0 | 10.0 |
| R4 | 10.0 | 10.0 | 10.0 | 10.0 | n/a | n/a | n/a | n/a | 10.0 | 10.0 | 10.0 | 10.0 |
| R1 | 147.0 | 147.0 | 138.0 | 144.0 | 151.0 | 131.0 | 129.0 | 137.0 | 151.0 | 161.0 | 122.0 | 144.7 |
| R2 | 213.0 | 189.0 | 216.0 | 206.0 | 223.0 | 270.0 | 206.0 | 233.0 | 193.0 | 157.0 | 256.0 | 202.0 |
| R3 | 323.0 | 318.0 | 337.0 | 326.0 | 343.0 | 280.0 | 299.0 | 307.3 | 254.0 | 272.0 | 355.0 | 293.7 |
| R4 | 222.0 | 231.0 | 265.0 | 239.3 | n/a | n/a | n/a | n/a | 227.0 | 238.0 | 261.0 | 242.0 |

**Figure 1 legend:**
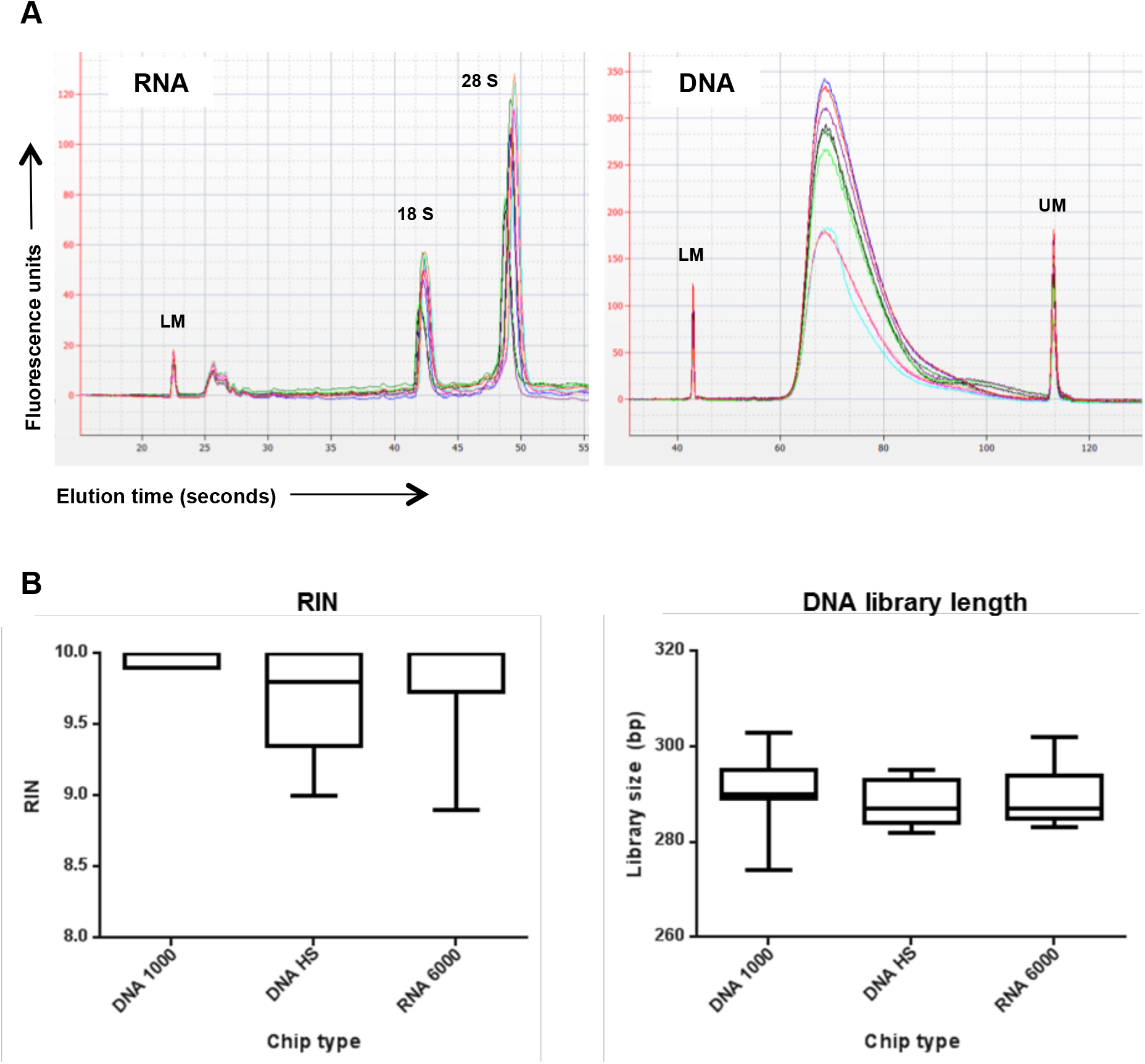
Inter- and intra-chip consistency in the measurement of key sample parameters. A. Overlaid electropherogram traces of all RNA and DNA Bioanalyzer runs. Four RNA or DNA samples were run in triplicate on an RNA 6000 Nano, DNA High Sensitivity (HS) and DNA 1000 chip, according to the RNA 6000 Nano or DNA HS kit protocol, respectively. Note that the lower fluorescence for the DNA 1000 chip did not affect calculations of sample concentration or length. B. RNA integrity number (RIN) of the four RNA samples and library size of the four RNA-seq libraries were determined on each chip. UM = Upper marker; LM = Lower marker peak.

The size distribution and concentration of four DNA samples (RNA-seq libraries; D1, D2, D3, and D4) were assessed using the DNA HS reagents and protocol, on RNA 6000 Nano, DNA HS and DNA 1000 chips; again, the sticker displaying the chip layout was disregarded. All chips were run on the Bioanalyzer using the DNA HS assay. Manual integration was used to label the prominent peak for each RNA-seq library, enabling library length and concentration to be calculated. The average size estimated across all Bioanalyzer chips was 290 bp (SD 6 bp) **(Table 2, and Fig. 1)**, with a 40 bp range across all samples (inter-chip library length: ANOVA p = 0.69). DNA concentration showed a 10-20 % variation in concentration across chips **(Table 2).** Accordingly, care should be taken when quantifying next-generation sequencing libraries using the Agilent Bioanalyzer; however, the recommendation is to use quantitative PCR (7).

**Table 2 legend:** DNA data. Four DNA samples (RNA-seq libraries) were run in triplicate according to the DNA High Sensitivity (HS) protocol, on a DNA 1000, DNA HS and RNA 6000 Nano chip. Manual integration was used to delete artefacts and select the desired peak; library length (upper) and concentration data (lower) (ng/μl) were determined by the software. The DNA HS protocol uses 11 sample wells, therefore D4 was loaded in duplicate. M = median library length or mean concentration.

| DNA Sample | DNA 1000 |  |  |  | DNA HS |  |  |  | RNA Nano |  |  |  |
| --- | --- | --- | --- | --- | --- | --- | --- | --- | --- | --- | --- | --- |
|  | 1 | 2 | 3 | Md | 1 | 2 | 3 | Md | 1 | 2 | 3 | Md |
| D1 | 274 | 284 | 291 | 284 | 287 | 285 | 282 | 285 | 286 | 285 | 283 | 285 |
| D2 | 303 | 294 | 297 | 297 | 294 | 292 | 295 | 294 | 295 | 294 | 290 | 294 |
| D3 | 290 | 290 | 290 | 290 | 282 | 285 | 284 | 284 | 286 | 285 | 287 | 286 |
| D4 | 295 | 289 | n/a | 292 | 293 | 293 | n/a | 293 | 302 | 293 | n/a | 298 |
| D1 | 2.4 | 2.1 | 2.1 | 2.2 | 1.9 | 2.0 | 2.0 | 2.0 | 2.7 | 2.6 | 2.5 | 2.6 |
| D2 | 5.3 | 6.3 | 6.0 | 5.9 | 5.3 | 5.7 | 5.3 | 5.4 | 5.7 | 6.6 | 6.1 | 6.1 |
| D3 | 2.0 | 1.9 | 1.9 | 1.9 | 1.8 | 1.7 | 1.8 | 1.8 | 2.3 | 2.5 | 2.3 | 2.4 |
| D4 | 4.6 | 6.2 | n/a | 5.4 | 4.7 | 5.3 | n/a | 5.0 | 4.5 | 6.2 | n/a | 5.3 |

Four DNA samples (RNA-seq libraries) were run in triplicate according to the DNA High Sensitivity (HS) protocol, on each chip. Manual integration was used to delete artefacts and select the desired peak; library length (bp) (upper) and concentration data (lower) (ng/μl) were determined by the software. The DNA HS protocol uses 11 sample wells, therefore D4 was loaded in duplicate. Md = median.

To assess assay reproducibility, which has been previously described by others (8,9), a pool of the RNA samples described above, or a single DNA sample (RNA-seq library pool, SLX-10140) were loaded as technical replicates in each well of duplicate chips, for all three chip types tested. Library length and RIN were consistent between all chips (Fig. 2). RNA and DNA concentration data were significantly different between chip types, although it was apparent that this was due to inter-chip variability and not due to chip type (Fig. 2 and Supplementary data).

**Figure 2 legend:**
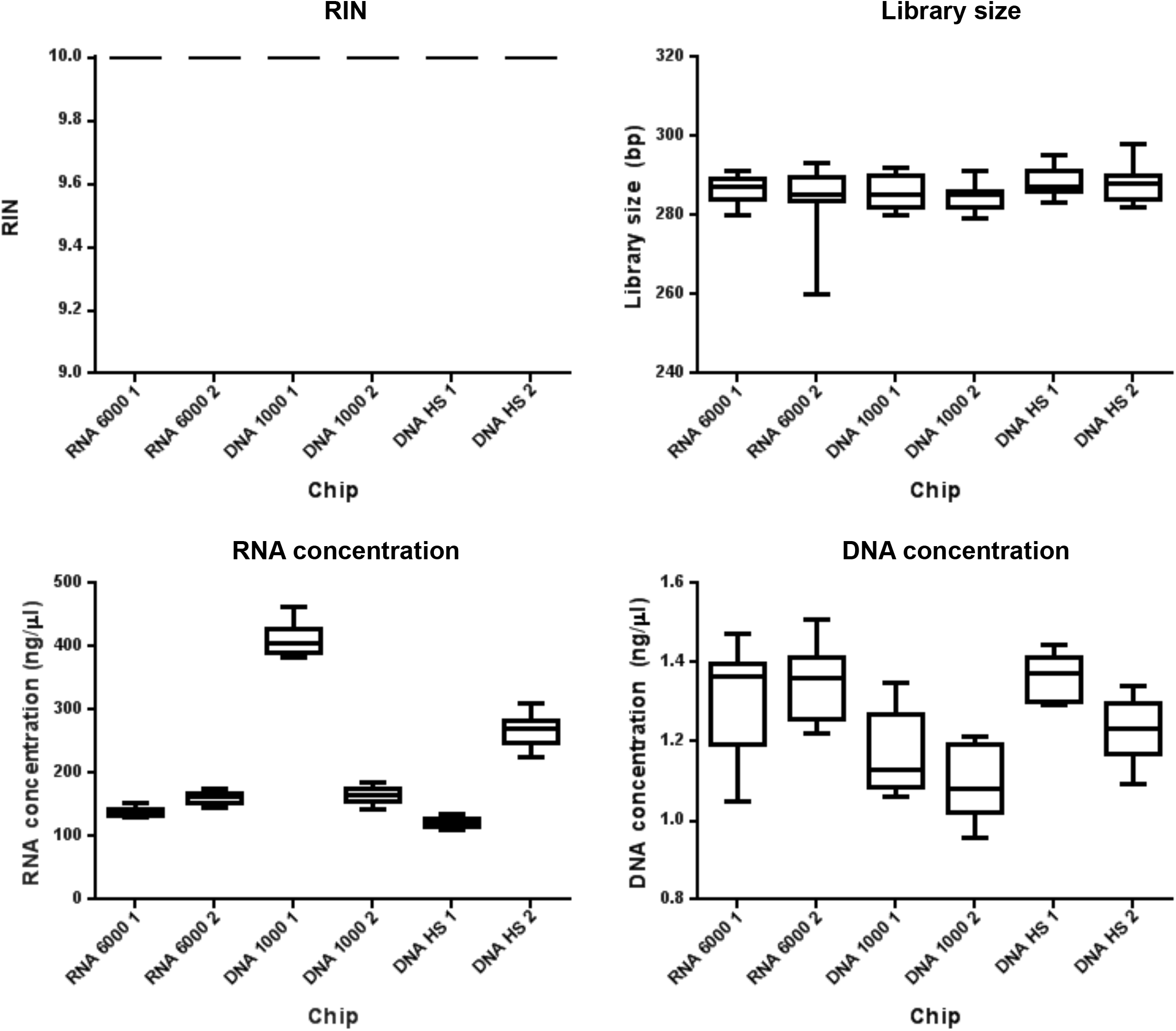
Reproducibility of chip sample metrics. A technical replicate of RNA or DNA was loaded onto duplicates of each chip type, and the RNA 6000 Nano or DNA HS kit protocols, respectively, were followed. Tabulated data and results of statistical tests are given in the Supplementary material.

We determined the accuracy of our Bioanalyzer concentration measurements by comparison to NanoDrop UV spectrophotometry (Nanodrop, DE, USA), Qubit (Thermo Fisher Scientific, MA, USA) and Kapa qPCR (KAPA BioSystems, South Africa) measurements (Qubit and qPCR are the recommended quantification methods for RNA and RNA-seq libraries, respectively). The Bioanalyzer concentration measurement of the RNA pool was less precise (211 ng/μl, SD 102.89 ng/μl) than Qubit (146 ng/μl, SD 6.24 ng/μl). The Bioanalyzer underestimated the concentration of our RNA-seq library (1.25 ng/μl) when compared to qPCR (11.17 ng/μl) (Supplementary data). Our data support the finding that Bioanalyzer quantification is more variable than other methods (9).

Our data confirm that, provided the assay-specific protocol is followed, the Bioanalyzer chip type used is irrelevant for RIN and DNA size estimation, i.e. qualitative analysis of RNA and RNA-seq libraries. It is important that the chip sticker is not used as a guide. Qubit and qPCR are recommended for RNA and RNA-seq library concentration measurements; the variability in RNA and DNA concentration measurements identified between Bioanalyzer runs supports this. These findings will be applicable to an extensive number of research environments in which the Agilent 2100 Bioanalyzer is used for the assessment of nucleic acid samples; in particular, those facilities which employ more than one type of kit and are high consumers of Bioanalyzer reagents.

## Acknowledgements

We thank all members of the Genomics Core team past and present for useful discussions; Sarah Vowler for help with statistical tests; and Cancer Research UK and the University of Cambridge for funding the Genomics Core facilities through the Cambridge Institute grant.

